# COVID-19 mRNA third dose induces a unique hybrid immunity-like antibody response

**DOI:** 10.1101/2022.05.09.491201

**Authors:** Emanuele Andreano, Ida Paciello, Giulio Pierleoni, Giulia Piccini, Valentina Abbiento, Giada Antonelli, Piero Pileri, Noemi Manganaro, Elisa Pantano, Giuseppe Maccari, Silvia Marchese, Lorena Donnici, Linda Benincasa, Ginevra Giglioli, Margherita Leonardi, Concetta De Santi, Massimiliano Fabbiani, Ilaria Rancan, Mario Tumbarello, Francesca Montagnani, Claudia Sala, Duccio Medini, Raffaele De Francesco, Emanuele Montomoli, Rino Rappuoli

## Abstract

The continuous evolution of SARS-CoV-2 generated highly mutated variants, like omicron BA.1 and BA.2, able to escape natural and vaccine-induced primary immunity^1,2^. The administration of a third dose of mRNA vaccines induces a secondary response with increased protection. We investigated, at single-cell level, the longitudinal evolution of the neutralizing antibody response in four donors after three mRNA doses^3^. A total of 4,100 spike protein specific memory B cells were single cell sorted and 350 neutralizing antibodies were identified. The third dose increased the antibody neutralization potency and breadth against all SARS-CoV-2 variants of concern as previously observed with hybrid immunity^3^. However, the B cell repertoire that stands behind the response is dramatically different. The increased neutralizing response was largely due to the expansion of B cell germlines poorly represented after two doses, and the reduction of germlines predominant after primary immunization such as IGHV3-53;IGHJ6-1 and IGHV3-66;IGHJ4-1. Divergently to hybrid immunity, cross-protection after a third dose was mainly guided by Class 1/2 antibodies encoded by IGHV1-58;IGHJ3-1 and IGHV1-69;IGHJ4-1 germlines. The IGHV2-5;IGHJ3-1 germline, which induced broadly cross-reactive Class 3 antibodies after infection or viral vector vaccination, was not induced by a third mRNA dose. Our data show that while neutralizing breadth and potency can be improved by different immunization regimens, each of them has a unique molecular signature which should be considered while designing novel vaccines and immunization strategies.

## INTRODUCTION

The emergence of SARS-CoV-2 variants of concern (VoCs) able to escape vaccine immunity elicited by the spike (S) protein of the original virus isolated in Wuhan, China, have decreased the impact of vaccination^4,5^. As a consequence, the COVID-19 pandemic continues to impact the health, the economy and the freedom of people worldwide in spite of the several billion doses of vaccines already deployed. This scenario raises new important questions about the use of the existing vaccines and the immunity they can provide to tackle current and future variants. Therefore, it became of utmost importance to understand the nature and the quality of the immune response elicited by the different vaccines used worldwide. In our previous study we analyzed at single cell level the immune response induced by two doses of the BNT162b2 mRNA vaccine in naïve people and in people that had been previously infected by the SARS-CoV-2 virus^3^. In this study we analyzed at single cell level the longitudinal B cell and neutralizing antibody response of the same naïve people after a third immunization. We found that, while the overall immune response after a third dose is similar to that observed in the hybrid immunity of vaccinated people previously infected by the virus, the B cell repertoire that stands behind the response is dramatically different.

## RESULTS

### B cells response after third dose

To evaluate the longitudinal evolution of the neutralizing antibody response, four seronegative donors that participated in our previous study after two doses of BNT162b2 mRNA vaccine^3^, were re-enrolled after receiving a third dose. None of the subjects were exposed to SARS-CoV-2 infection between the second and third vaccination dose. Data after the third dose (seronegative 3^rd^ dose; SN3) described in this study were compared to those obtained from the same subjects after the second dose (seronegative 2^nd^ dose; SN2) and to those of subjects with hybrid immunity (seropositive 2^nd^ dose; SP2) previously described^3^. Three subjects received the BNT162b2 (VAC-001, VAC-002 and VAC-008) vaccine, while one subject (VAC-010) received the mRNA-1273 vaccine. Blood collection occurred at an average of 58 days post third vaccination dose. Subject details are summarized in **Extended Data Table 1**. The frequency of CD19^+^CD27^+^IgD^-^IgM^-^ memory B cell (MBCs) specific for the S protein^+^ was 6.18-fold higher in SN3 compared to SN2 (**Fig. 1a-b**). This was also 2.26-fold higher than that previously observed in subjects with SARS-CoV-2 infection and subsequent vaccination with two doses of the BNT162b2 mRNA vaccine (SP2)^3^. No major differences between the two groups were observed in the CD19^+^CD27^+^IgD^-^IgM^-^/IgM^+^ memory B cell (MBCs) compartments (**Extended Data Fig. 1a-d**). In addition, SN3 showed higher binding to the S protein, receptor binding domain (RBD) and N-terminal domain (NTD), and a 5.82-fold higher neutralization activity against the original Wuhan SARS-CoV-2 virus compared to SN2 (**Fig. 1c-d; Extended Data Fig. 1e-h**).

**Fig. 1.**
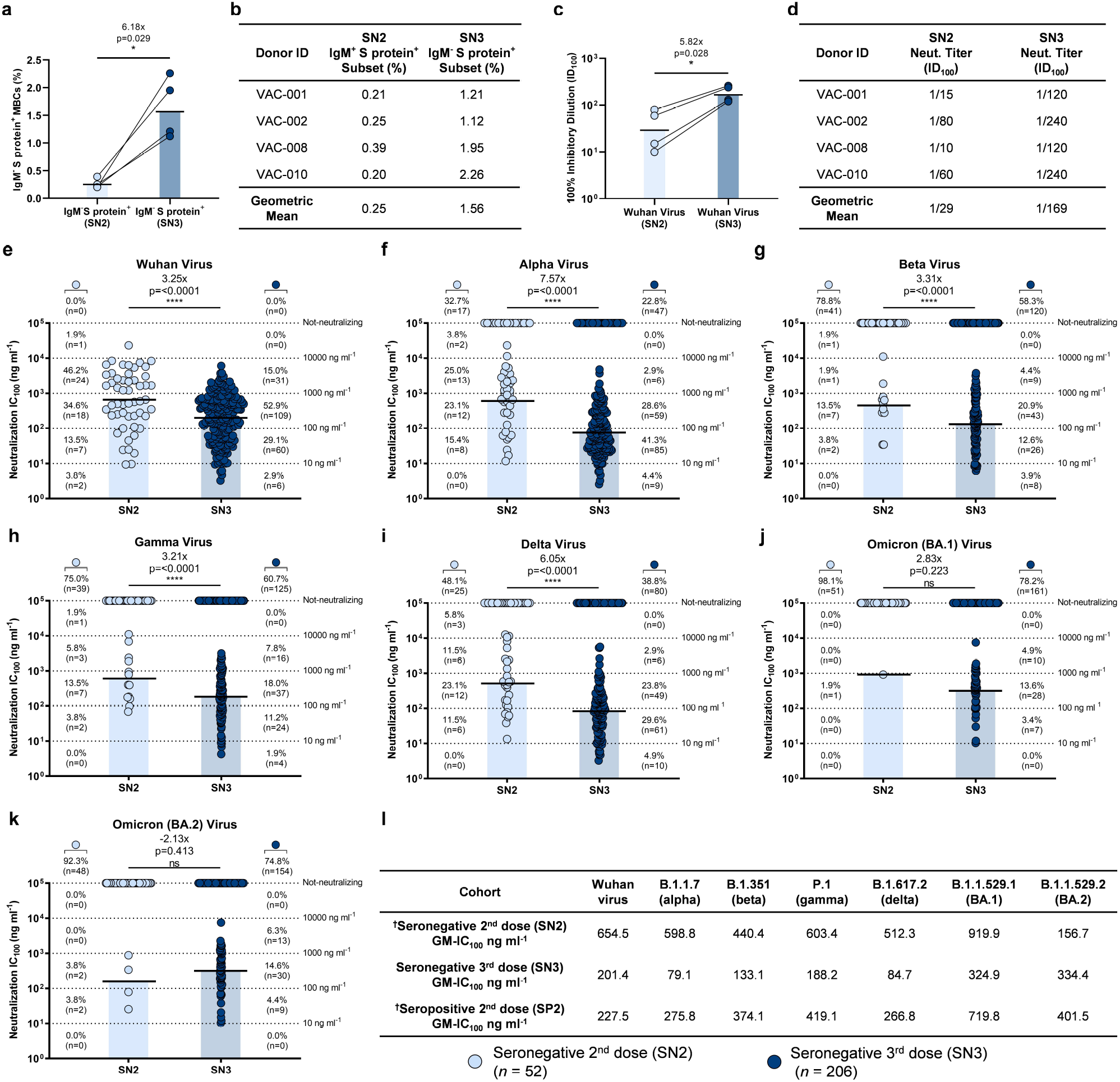
Potency and breadth of neutralization of nAbs against SARS-CoV-2 and VoCs. **a**, The graph shows the frequency of CD19^+^CD27^+^IgD^-^IgM^-^ able to bind the SARS-CoV-2 S protein trimer (S protein^+^) in SN2 and SN3 donors (*n* = 4/group). Black line and bars denote the geometric mean. **b**, The table summarizes the frequencies of the B cell populations in SN2 and SN3. **c**, The graph shows the neutralizing activity of plasma samples against the original Wuhan SARS-CoV-2 virus. Technical duplicates were performed for each experiment. **d**, The table summarizes the 100% inhibitory dilution (ID_100_) of each COVID-19 vaccinee and the geometric mean for SN2 and SN3 (*n* = 4/group). **e-k**, Scatter dot charts show the neutralization potency, reported as IC_100_ (ng ml^-1^), of nAbs tested against the original Wuhan SARS-CoV-2 virus (**e**) and the alpha (**f**), beta (**g**), gamma (**h**), delta (**i**), omicron BA.1 (**j**) and omicron BA.2 (**k**) VoCs. The number and percentage of nAbs from SN2 vs SN3, fold-change and statistical significance are denoted on each graph. **l**, The table shows the IC_100_ geometric mean (GM) of all nAbs pulled together from SN2, SN3 and SP2 against all SARS-CoV-2 viruses tested. “†” indicates previously published data^3^. Technical duplicates were performed for each experiment. A nonparametric Mann–Whitney t test was used to evaluate statistical significances between groups. Two-tailed p-value significances are shown as *p < 0.05, **p < 0.01, ***p < 0.001, and ****p < 0.0001.

### Boosting antibody potency and breadth

To evaluate the longitudinal B cell response, antigen specific class-switched MBCs (CD19^+^CD27^+^IgD^-^IgM^-^) were single cell sorted using as bait the Wuhan prefusion SARS-CoV-2 S protein trimeric antigen which was encoded by the mRNA vaccine. Sorted cells were incubated for two weeks to naturally release human monoclonal antibodies (mAbs) into the supernatant. A total of 4,100 S protein^+^ MBCs were sorted and 2,436 (59.4%) produced mAbs able to recognize the S protein prefusion trimer in ELISA. Of these, 350 neutralized the original Wuhan live SARS-CoV-2 virus when tested at a single point dilution (1:10) by cytopathic effect-based microneutralization assay (CPE-MN). Overall, the fraction of S protein-specific B cells producing neutralizing antibodies (nAbs) was 14.4% which is 2.2-fold higher than what observed in SN2 and comparable to what observed for SP2 dose vaccinees (14.8%) in our previous study (**Extended Data Fig. 2a-b**; **Extended Data Table 2**)^3^. To better characterize identified nAbs, we tried to express all 350 as immunoglobulin G1 (IgG1), and we were able to recover and express 206 of them. Binding by ELISA to RBD and NTD of the original Wuhan SARS-CoV-2 S had similar frequency in SN2 and SN3, while we observed a reduction of S protein trimer specific antibodies after a third booster dose, a trend similar to what observed in SP2 (**Extended Data Fig. 2c**). No S2 domain binding nAbs were identified. Finally, we evaluated the neutralization potency against the original Wuhan virus, the alpha (B.1.1.7), beta (B.1.351), gamma (P.1) delta (1.617.2), omicron BA.1 (B.1.1.529.1) and omicron BA.2 (B.1.1.529.2) and compared it with SN2 (**Fig. 1e-l**). To increase the power of the analyses, in Fig.1 we included all 52 nAbs isolated from the five seronegative donors enrolled in our previous study^3^. Overall, nAbs isolated in SN3 were higher in frequency and potency against all tested viruses compared to SN2 and showed a significantly higher 100% inhibitory concentration (IC_100_), except for BA.1 and BA.2 VoCs, where SN2 did not have enough nAbs to make a meaningful comparison (**Fig. 1e-l**). The IC_100_ geometric mean (GM-IC_100_) in SN3 was 3.25, 7.57-, 3.31-, 3.21-, 5.70- and 2.83-fold lower compared to SN2 for the Wuhan virus, the alpha, beta, gamma, delta and BA.1 VoCs respectively (**Fig. 1; Extended Data Fig. 3**). Only for BA.2 a higher GM-IC_100_ was observed in SN2 compared to SN3. Interestingly, SN3 also showed a higher neutralization potency compared to SP2 against all tested viruses with a 1.13-, 3.49-, 2.81, 2.23-, 3.15-, 2.22-, and 1.20-fold lower GM-IC_100_ (**Fig. 1l; Extended Data Fig. 3**). In addition, we observed that nAbs induced following a third booster dose retained a high ability to cross-neutralize all tested SARS-CoV-2 VoCs. Indeed, while 67.3, 21.1, 25.0, 51.9, 1.9 and 7.7% of nAbs maintained the ability to neutralize alpha, beta, gamma, delta, omicron BA.1 and omicron BA.2 respectively in the SN2 group^3^, the same viruses were neutralized by 77.2, 41.7, 39.3, 61.2, 21.8 and 25.2% of nAbs in the SN3 group (**Fig. 1; Extended Data Table 2**). Finally, we assessed if a third booster dose would enhance the ability of nAbs to recognize and cross-neutralize the distantly related SARS-CoV-1 virus. Two of the four SN3 donors presented nAbs able to recognize SARS-CoV-1 (**Extended Data Fig. 4a**). However, only 2.4% (5/206) and 0.9% (2/206) of all nAbs were able to bind the SARS-CoV-1 S protein and neutralize this sarbecovirus respectively, showing a lower frequency than the one observed in SN2 and SP2 nAbs (**Extended Data Fig. 4a-b**)^3^.

### Antibody classes of cross-protection

To understand the epitope region recognized by our RBD targeting-antibodies (*n* = 154) after a third vaccination dose, we performed a competition assay with three known antibodies. As previously described, we used the Class 1/2 antibody J08^6^, the Class 3 antibody S309^7^, and the Class 4 antibody CR3022^8^ to map the epitope regions targeted by our RBD-binding nAbs^3^. The most abundant class of nAbs targeted the Class 1/2 epitope region (98/154; 63.6%), followed by Class 3 targeting nAbs (30/154; 19.5%) and nAbs that recognize the Class 4 region (3/154; 1.9%). The remaining 23 (14.9%) nAbs did not compete with any of the three antibodies used in our competition assay. As shown in **Fig. 2a**, in SN3 we observed a marked increase in Class 3 and 4 nAbs compared to SN2 and a similar distribution to SP2. When we compared the GM-IC_100_ of classes of nAbs in the SN3 with the SN2 and SP2 groups, we observed similar neutralization potency when tested against the SARS-CoV-2 virus originally isolated in Wuhan, China (**Extended Data Fig. 5**). Following, we aimed to understand which classes of RBD (*n* = 154) and NTD (*n* = 43) targeting nAbs in SN3 were mainly responsible for the cross-protection against the SARS-CoV-2 VoCs. Overall, we observed that Class 1/2 nAbs are the most abundant family of antibodies against all tested variants, followed by Class 3, Not-competing, NTD and Class 4 nAbs (**Fig. 2b-c**). Finally, we compared the functional antibody response between SN3 and SP2 against the highly mutated omicron BA.1 and BA.2 viruses. Interestingly, we observed that a third mRNA booster dose increases neutralization potency and evasion resistance of Class 1/2 nAbs against both omicron BA.1 and BA.2 compared to SP2. The opposite trend was observed for Class 3 nAbs, which had a lower frequency of cross-protection in SN3 compared to SP2 (**Fig. 2d-e**).

**Fig. 2.**
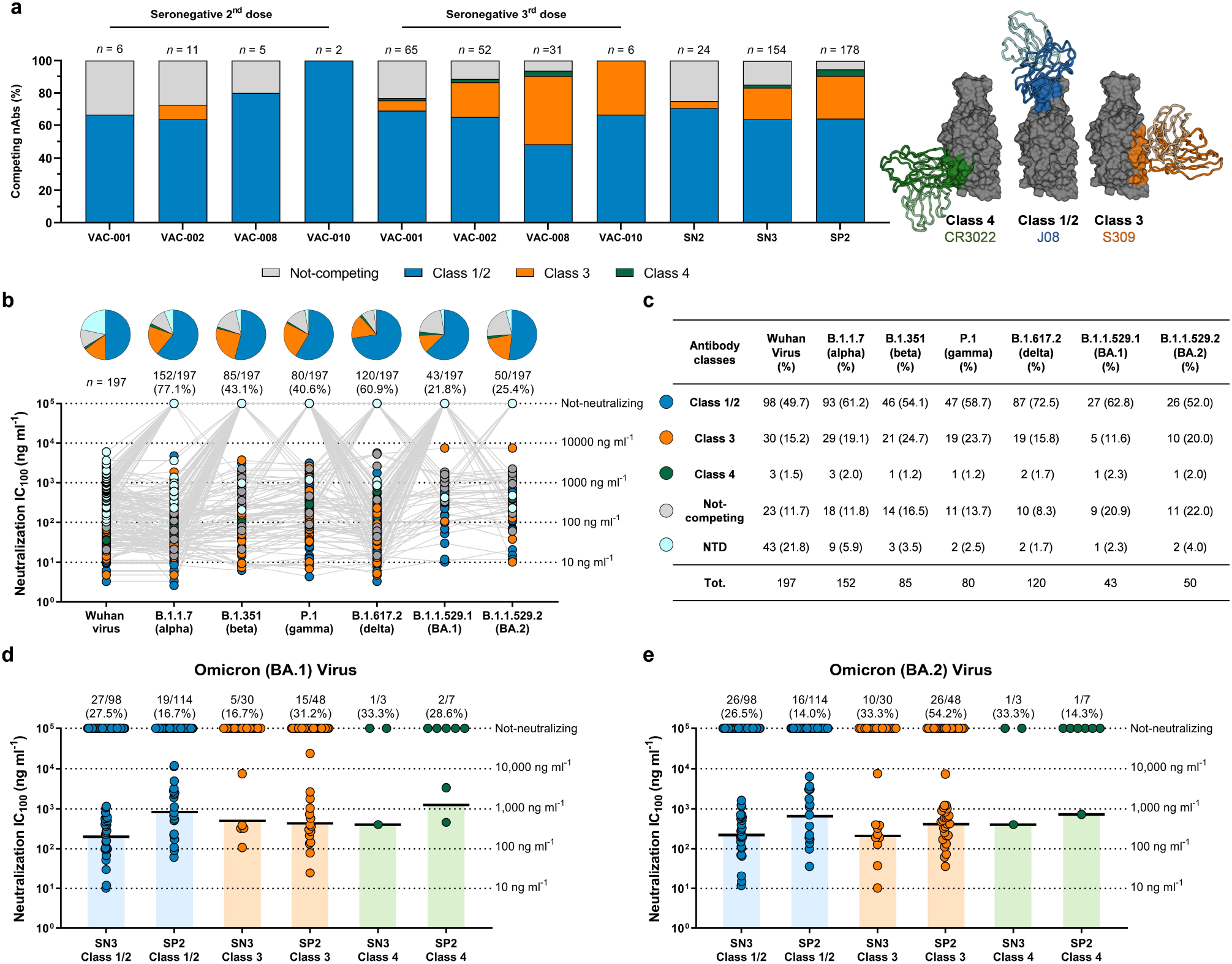
Distribution of SN3 nAbs against SARS-CoV-2 VoCs. **a**, The bar graph shows the epitope regions recognized by RBD-binding nAbs. Class 1/2, Class 3, Class 4 and Not-competing nAbs are shown in light blue, orange, green and light gray respectively. The number (*n*) of antibodies tested per each donor is denoted on the graph. Right panel highlights the epitope recognized by Class 1/2 (J08), Class 3 (S309) and Class 4 (CR3022) antibodies used in our competition assay. **b**, Pie charts show the distribution of cross-protective nAbs based on their ability to bind Class 1/2, Class 3 and Class 4 regions on the RBD, as well as not-competing nAbs (gray) and NTD-targeting nAbs (cyan). Dot charts show the neutralization potency, reported as IC_100_ (ng ml^-1^), of nAbs against the Wuhan virus and alpha, beta, gamma, delta, omicron BA.1 and omicron BA.2 SRS-CoV-2 VoCs observed in the SN3 group. The number and percentage of nAbs are denoted on each graph. **c**, the table summarizes number and percentages of Class 1/2, Class 3, Class 4, not-competing and NTD-targeting nAbs for each tested variant. **d-e**, dot charts compare the distribution of nAbs between SN3 and SP2 groups against omicron BA.1 (**d**) and BA.2 (**e**). The number, percentage and GM-IC_100_ (black lines and colored bars) of nAbs are denoted on each graphs.

### Antibody gene repertoire

We then interrogated the functional antibody repertoire. Initially, we analyzed all immunoglobulin heavy chain sequences retrieved from the three different groups (SN2 *n* = 58; SN3 *n* = 288; SP2 *n* = 278), and their respective V-J gene rearrangements (IGHV;IGHJ)^3^. Interestingly, SN2 and SN3 share only 20.2% percent (23/114) of SN3 IGHV;IGHJ rearrangements, while SN3 and SP2 share up to 45.6% (52/114). In addition, we observed that the frequency of antibodies encoded by predominant rearrangements induced by 2 vaccination doses (i.e. IGHV3-30;IGHJ6-1, IGHV3-33;IGHJ4-1, IGHV3-53;IGHJ6-1 and IGHV3-66;IGHJ4-1)^3,6,9,10^ were reduced after a third booster dose, while we observed an expansion of the antibody germlines IGHV1-58;IGHJ3-1 and IGHV1-69;IGHJ4-1 which were previously found to be predominant in SP2^3,11^ (**Fig. 3a; Extended Data Fig. 6**). These latter germlines previously showed high level of cross-neutralization activity against SARS-CoV-2 VoCs^12-14^. In addition, we observed in one donor (VAC-001) an important expansion of the germline IGHV1-46;IGHJ6-1 after receiving a 3^rd^ booster dose (**Fig. 3a; Extended Data Fig. 6**). Conversely to what found in SP2^3^, we did not observe the expansion of the Class 3 targeting antibody germline IGHV2-5;IGHJ4-1, which so far has been observed only in previously infected vaccinees or subjects immunized with adenoviral vectors^3,15,16^. Following, we aimed to identify expanded clonal families within the same four donors after a second and third vaccination dose. Sequences (SN2 *n* = 43; SN3 *n* = 288) were clustered by binning the clones to their inferred germlines (centroids) and according to 80% nucleotide sequence in the heavy complementary determining region 3 (CDRH3). Clusters were defined as antibody families including at least five or more members as previously described^17^. Of the 331 sequences, 226 (68.3%) were orphans (i.e. did not cluster with other sequences), and only in six cases sequences from SN2 and SN3 were binned to the same centroid (**Fig. 3b**). Only five clusters were identified, three of which were composed by antibodies belonging exclusively from the SN3 group. Of these clusters, two were formed by antibody sequences shared between the SN2 and SN3 groups. The smallest cluster was composed by 11 antibody members encoded from the IGHV1-58;IGHJ3-1 germline, while the biggest cluster, composed by 18 antibody members, were encoded by the IGHV3-48/IGHV3-53/IGHV3-66 germlines (**Fig. 3b**). Finally, we evaluated the V-gene mutation levels, neutralization potency and breadth in nAbs encoded by reduced and expanded germlines following a third vaccination dose. Our data showed that IGHV3-53;IGHJ6-1 and IGHV3-66;IGHJ4-1 germlines, expanded in SN2 and reduced in SN3, have a similar level of V gene somatic mutations, are poorly cross-reactive especially against omicron BA.1 and BA.2, and show medium neutralization potency (IC_100_ mainly between 100 and 1,000 ng ml^-1^). Differently, the antibody germlines IGHV1-58;IGHJ3-1 and IGHV1-69;IGHJ4-1, mainly expanded in SN3 and poorly represented in SN2, show between 5 and 15-fold higher V gene somatic mutation levels, higher neutralization potency and breadth against all SARS-CoV-2 VoCs (**Fig. 3c-f**).

**Fig. 3.**
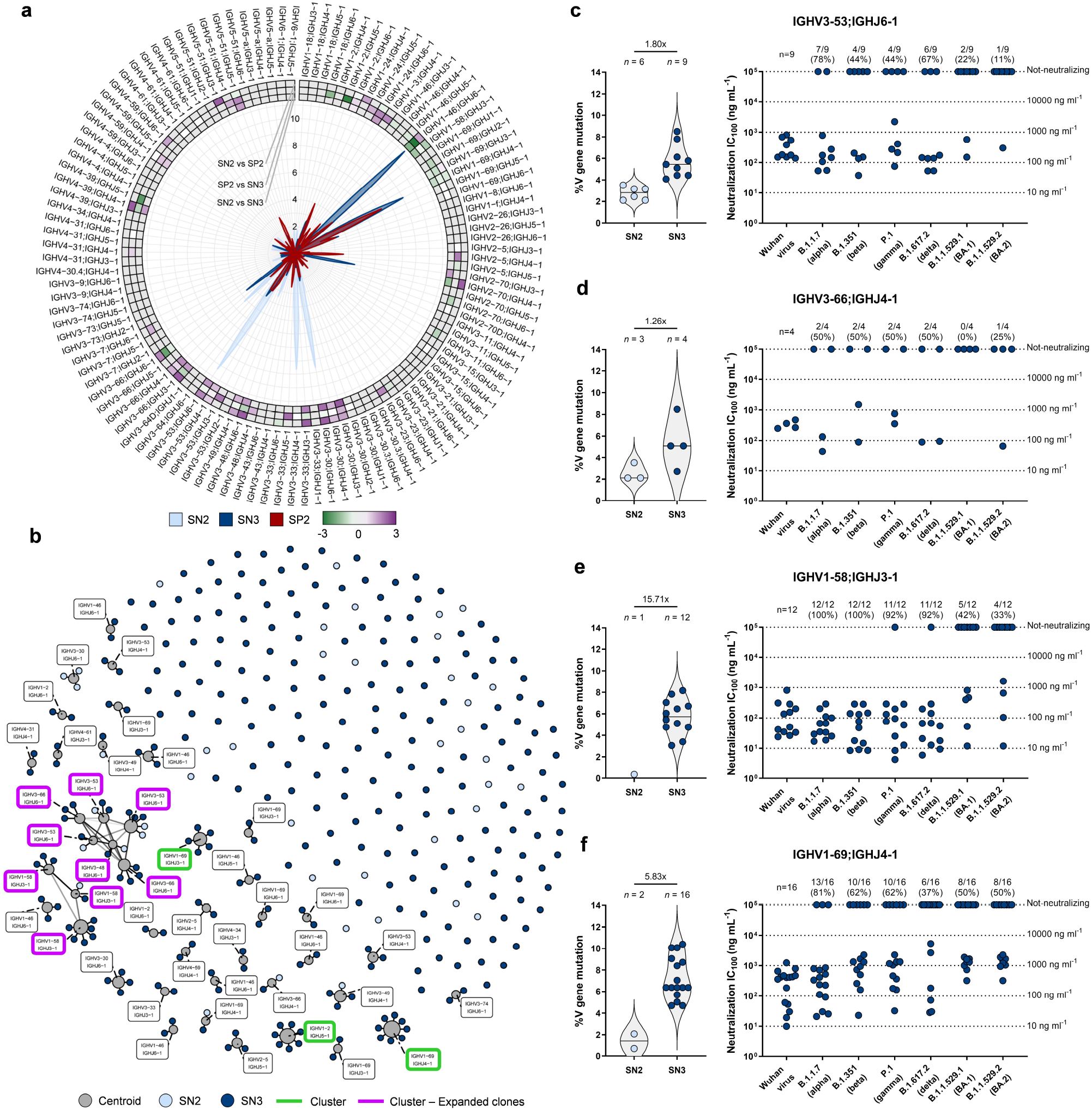
Repertoire analyses and characterization of predominant antibody germlines. **a**, Radar plot shows the distribution of IGHV;IGHJ germlines among the three different groups. SN2, SN3 and SP2 are shown in light blue, dark blue and red respectively. Heatmap represents the Log2 fold change (FC) among groups. **b**, Network plot shows the clonally-expanded antibody families in SN2 and SN3. Centroids, nAbs from SN2 and SN3 groups are shown in gray, light blue and dark blue respectively. Clusters and expanded clones are highlighted in bright green and bright purple respectively. **c-f**, The graphs show the V gene somatic mutation frequency (left panel) and neutralization potency (IC_100_; right panel) of IGHV3-53;IGHJ6-1 (**c**), IGHV3-66;IGHJ4-1 (**d**), IGHV1-58;IGHJ3-1 (**e**), and IGHV1-69;IGHJ4-1 (**f**) gene derived nAbs, against the original SARS-CoV-2 virus first detected in Wuhan, China, and all SARS-CoV-2 VoCs. The number and percentage of nAbs analyzed are denoted on each graph.

## DISCUSSION

In agreement with other longitudinal studies^9,10,18^, we found that a third dose of mRNA vaccine induces an immune response similar to the hybrid immunity observed in people vaccinated after SARS-CoV-2 infection. This antibody response is characterized by a small increase in S protein binding antibodies, a strong increase in neutralizing potency and a considerable increase in antibodies able to cross-neutralize emerging variants, including omicron BA.1 and BA.2. The increased potency and breadth are due to a significant expansion of S protein specific MBCs, which is even higher than that observed in subjects with hybrid immunity, and by a strong increase of V gene somatic mutations. What is new in our study is that the increased potency and breadth observed after a third booster dose was due mostly to Class 1/2 nAbs, while Class 3 antibodies had a lower frequency and breadth compared to subjects with hybrid immunity. In addition, we found that a third mRNA dose did not induce a strong response against the distantly related SARS-CoV-1, suggesting that additional doses of homologous vaccines against SARS-CoV-2 will focus the antibody response against this virus instead of broadening cross-protection to other coronaviruses. Another unique observation of this study is that the increased neutralization potency and breadth is not due to a linear evolution of the B cells producing nAbs after two vaccine doses but is due mostly to the expansion of new B cells which were not detected after primary immunization. Indeed, the secondary response induced by a third vaccine dose did not derive from expanded B cell clones, but it was dominated by singlets that constituted 68% of the entire repertoire. In addition, the germlines IGHV3-53;IGHJ6-1/IGHV3-66;IGHJ4-1 which dominated the neutralizing response in donors infected with the original Wuhan virus and in subjects immunized with two doses of mRNA vaccines^3,6,19-21^, decreased in frequency and did not improve in potency or cross-neutralization after a third dose. Conversely, the antibody germlines IGHV1-58;IGHJ3-1 and IGHV1-69;IGHJ4-1 became largely responsible for the improved potency and cross-neutralization observed after a third dose. Interestingly, the IGHV1-58 germline predominant in this group was previously shown to recognize a “supersite” on the S protein surface and to be expanded following beta or omicron breakthrough infection in vaccinated individuals^12,14,22^. A unique observation of this work is that the highly cross-reactive germline IGHV2-5;IGHJ4-1, found in subjects with hybrid immunity and in people vaccinated with viral vectors^3,16^, is absent after a third mRNA vaccine, suggesting that the induction of this germline may require endogenous production of the S protein. The lack of expansion of the IGHV2-5;IGHJ4-1 germline after three mRNA vaccine doses could partially explain the above mentioned loss of cross-reactivity by Class 3 nAbs. Overall, our study provides a high-resolution picture of the functional and genetic properties of a third mRNA vaccination and, despite we observed important similarities with the hybrid immunity, unravels features of the antibody response uniquely produced after a third mRNA vaccine dose. These observations are very important and should be considered while designing new vaccines and implementing vaccination regimens for booster doses, especially in low-middle income countries where less than 20% of people received at least one dose of vaccines^23^.

## METHODS

### Enrollment of COVID-19 vaccinees and human sample collection

This work results from a collaboration with the Azienda Ospedaliera Universitaria Senese, Siena (IT) that provided samples from COVID-19 vaccinated donors, of both sexes, who gave their written consent. The study was approved by the Comitato Etico di Area Vasta Sud Est (CEAVSE) ethics committees (Parere 17065 in Siena) and conducted according to good clinical practice in accordance with the declaration of Helsinki (European Council 2001, US Code of Federal Regulations, ICH 1997). This study was unblinded and not randomized. No statistical methods were used to predetermine sample size.

### Single cell sorting of SARS-CoV-2 S protein^+^ memory B cells from COVID-19 vaccinees

Peripheral blood mononuclear cells (PBMCs) and single cell sorting strategy were performed as previously described^3,6^. Briefly, PBMC were isolated from heparin-treated whole blood by density gradient centrifugation (Ficoll-Paque™ PREMIUM, Sigma-Aldrich) and stained with Live/Dead Fixable Aqua (Invitrogen; Thermo Scientific) diluted 1:500. After 20 min incubation cells were saturated with 20% normal rabbit serum (Life technologies) for 20 min at 4°C and then stained with SARS-CoV-2 S-protein labeled with Strep-Tactin®XT DY-488 (iba-lifesciences cat# 2-1562-050) for 30 min at 4°C. After incubation the following staining mix was used CD19 V421 (BD cat# 562440, 1:320), IgM PerCP-Cy5.5 (BD cat# 561285, 1:50), CD27 PE (BD cat# 340425, 1:30), IgD-A700 (BD cat# 561302, 1:15), CD3 PE-Cy7 (BioLegend cat# 300420, 1:100), CD14 PE-Cy7 (BioLegend cat# 301814, 1:320), CD56 PE-Cy7 (BioLegend cat# 318318, 1:80) and cells were incubated at 4°C for additional 30 min. Stained MBCs were single cell-sorted with a BD FACS Aria III (BD Biosciences) into 384-well plates containing 3T3-CD40L feeder cells, IL-2 and IL-21 and incubated for 14 days as previously described^24^.

### ELISA assay with SARS-CoV-2 and SARS-CoV-1 S protein prefusion trimer

mAbs and plasma binding specificity against the S-protein trimer was detected by ELISA as previously described^3^. Briefly, 384-well plates (microplate clear, Greiner Bio-one) were coated with 3 µg/mL of streptavidin (Thermo Fisher) diluted in carbonate-bicarbonate buffer (E107, Bethyl laboratories) and incubated at RT overnight. The next day, plates were incubated 1 h at RT with 3 µg/mL of SARS-CoV-2 or SARS-CoV-1 S protein, and saturated with 50 µL/well of blocking buffer (phosphate-buffered saline, 1% BSA) for 1 h at 37°C. Following, 25 µL/well of mAbs or plasma samples, diluted 1:5 or 1:10 respectively in sample buffer (phosphate-buffered saline, 1% BSA, 0.05% Tween-20), were added serially diluted step dilution 1:2 and then incubated at 1 h at 37°C. Finally, 25 µL/well of alkaline phosphatase-conjugated goat antihuman IgG and IgA (Southern Biotech) diluted 1:2000 in sample buffer were added. S protein binding was detected using 25 µL/well of PNPP (p-nitrophenyl phosphate; Thermo Fisher) and the reaction was measured at a wavelength of 405 nm by the Varioskan Lux Reader (Thermo Fisher Scientific). After each incubation step, plates were washed three times with 100 µL/well of washing buffer (phosphate-buffered saline, 0.05% Tween-20). Sample buffer was used as a blank and the threshold for sample positivity was set at 2-fold the optical density (OD) of the blank. Technical duplicates were performed for mAbs and technical triplicates were performed for sera samples.

### ELISA assay with RBD, NTD and S2 subunits

mAbs identification and plasma screening of vaccinees against RBD, NTD or S2 SARS-CoV-2 protein were performed by ELISA as previously described^3^. Briefly, 3 µg/mL of RBD, NTD or S2 SARS-CoV-2 protein diluted in carbonate-bicarbonate buffer (E107, Bethyl laboratories) were coated in 384-well plates (microplate clear, Greiner Bio-one) and blocked with 50 µL/well of blocking buffer (phosphate-buffered saline, 1% BSA) for 1h at 37°C. After washing, plates were incubated 1 h at 37 °C with mAbs diluted 1:5 in samples buffer (phosphate-buffered saline, 1% BSA, 0.05% Tween-20) or with plasma at a starting dilution 1:10 and step diluted 1:2 in sample buffer. Anti-Human IgG –Peroxidase antibody (Fab specific) produced in goat (Sigma) diluted 1:45000 in sample buffer was then added and samples incubated for 1 h at 37°C. Plates were then washed, incubated with TMB substrate (Sigma) for 15 min before adding the stop solution (H_2_SO_4_ 0.2M). The OD values were identified using the Varioskan Lux Reader (Thermo Fisher Scientific) at 450 nm. Each condition was tested in triplicate and samples tested were considered positive if OD value was 2-fold the blank.

### Flow cytometry-based competition assay

To classify mAbs candidates on the basis of their interaction with Spike epitopes, we performed a flow cytometry-based competition assay as previously described^3,11^. Briefly, magnetic beads (Dynabeads His-Tag, Invitrogen) were coated with histidine tagged S protein according to the manufacturers’ instructions. Then, 20 µg/mL of coated S protein-beads were pre-incubated with unlabeled nAbs diluted 1:2 in PBS for 40 minutes at RT. After incubation, the mix Beads-antibody was washed with 100 µL of PBS-BSA 1%. Then, to analyze epitope competition, mAbs able to bind RBD Class 1/2, (J08), Class 3 (S309) and Class 4 (CR3022) were labeled with three different fluorophores (Alexa Fluor 647, 488 and 594) using Alexa Fluor NHS Ester kit (Thermo Scientific), were mixed and incubated with S-protein-beads. Following 40 minutes of incubation at RT, the mix Beads-antibodies was washed with PBS, resuspended in 150 µL of PBS-BSA 1% and analyzed using BD LSR II flow cytometer (Becton Dickinson). Beads with or without S-protein incubated with labeled antibodies mix were used as positive and negative control respectively. FACSDiva Software (version 9) was used for data acquisition and analysis was performed using FlowJo (version 10).

### SARS-CoV-2 authentic viruses neutralization assay

All SARS-CoV-2 authentic virus neutralization assays were performed in the biosafety level 3 (BSL3) laboratories at Toscana Life Sciences in Siena (Italy) and Vismederi Srl, Siena (Italy). BSL3 laboratories are approved by a Certified Biosafety Professional and are inspected every year by local authorities. To evaluate the neutralization activity of identified nAbs against SARS-CoV-2 and all VoCs and evaluate the breadth of neutralization of this antibody is a cytopathic effect-based microneutralization assay (CPE-MN) was performed^3,6^. Briefly, the CPE-based neutralization assay sees the co-incubation of the antibody with a SARS-CoV-2 viral solution containing 100 median Tissue Culture Infectious Dose (100 TCID_50_) of virus for 1 hour at 37°C, 5% CO_2_. The mixture was then added to the wells of a 96-well plate containing a sub-confluent Vero E6 cell monolayer. Plates were incubated for 3-4 days at 37°C in a humidified environment with 5% CO_2_, then examined for CPE by means of an inverted optical microscope by two independent operators. All nAbs were tested a starting dilution of 1:5 and the IC_100_ evaluated based on their initial concentration while plasma samples were tested starting from a 1:10 dilution. Both nAbs and plasma samples were then diluted step 1:2. Technical duplicates were performed for both nAbs and plasma samples. In each plate positive and negative control were used as previously described^5^.

### SARS-CoV-2 virus variants CPE-MN neutralization assay

The SARS-CoV-2 viruses used to perform the CPE-MN neutralization assay were the original Wuhan SARS-CoV-2 virus (SARS-CoV-2/INMI1-Isolate/2020/Italy: MT066156), SARS-CoV-2 B.1.1.7 (INMI GISAID accession number: EPI_ISL_736997), SARS-CoV-2 B.1.351 (EVAg Cod: 014V-04058), B.1.1.248 (EVAg CoD: 014V-04089) and B.1.617.2 (GISAID ID: EPI_ISL_2029113)^26^.

### HEK293TN-hACE2 cell line generation

HEK293TN-hACE2 cell line was generated by lentiviral transduction of HEK293TN (System Bioscience) cells as described in Notarbartolo S. et al^25^. Lentiviral vectors were produced following a standard procedure based on calcium phosphate co-transfection with 3^rd^ generation helper and transfer plasmids. The transfer vector pLENTI_hACE2_HygR was obtained by cloning of hACE2 from pcDNA3.1-hACE2 (Addgene #145033) into pLenti-CMV-GFP-Hygro (Addgene #17446). HEK293TN-hACE2 cells were maintained in DMEM, supplemented with 10% FBS, 1% glutamine, 1% penicillin/streptomycin and 250 μg/ml Hygromicin (GIBCO).

### Production of SARS-CoV-1 pseudoparticles

SARS-CoV-1 lentiviral pseudotype particles were generated as described in Conforti et al. for SARS-CoV-2^26^ by using the SARS-CoV1 SPIKE plasmid pcDNA3.3_CoV1_D28 (Addgene plasmid # 170447).

### SARS-CoV-1 neutralization assay

For neutralization assay, HEK293TN-hACE2 cells were plated in white 96-well plates in complete DMEM medium. 24h later, cells were infected with 0.1 MOI of SARS-CoV-1 pseudoparticles that were previously incubated with serial dilution of purified or not purified (cell supernatant) mAb. In particular, a 7-point dose-response curve (plus PBS as untreated control), was obtained by diluting mAb or supernatant respectively five-fold and three-fold. Thereafter, nAbs of each dose-response curve point was added to the medium containing SARS-CoV-1 pseudoparticles adjusted to contain 0.1 MOI. After incubation for 1h at 37°C, 50 µl of mAb/SARS-CoV-1 pseudoparticles mixture was added to each well and plates were incubated for 24h at 37°C. Each point was assayed in technical triplicates. After 24h of incubation cell infection was measured by luciferase assay using Bright-Glo™ Luciferase System (Promega) and Infinite F200 plate reader (Tecan) was used to read luminescence. Obtained relative light units (RLUs) were normalized to controls and dose response curve were generated by nonlinear regression curve fitting with GraphPad Prism to calculate Neutralization Dose 50 (ND_50_).

### Single cell RT-PCR and Ig gene amplification and transcriptionally active PCR expression

To express our nAbs as full-length IgG1, 5 µL of cell lysate from the original 384-cell sorting plate were used for reverse transcription polymerase chain reaction (RT-PCR), and two rounds of PCRs (PCRI and PCRII-nested) as previously described^3,6^. Obtained PCRII products will be used to recover the antibody heavy and light chain sequences, through Sanger sequencing, and for antibody cloning into expression vectors as previously described^27,28^. Transcriptionally active PCR (TAP) reaction was performed using 5 μL of Q5 polymerase (NEB), 5 μL of GC Enhancer (NEB), 5 μL of 5X buffer,10 mM dNTPs, 0.125 µL of forward/reverse primers and 3 μL of ligation product, using the following cycles: 98°/2’, 35 cycles 98°/10’’, 61°/20’’, 72°/1’ and 72°/5’. TAP products were purified under the same PCRII conditions, quantified by Qubit Fluorometric Quantitation assay (Invitrogen) and used for transient transfection in Expi293F cell line following manufacturing instructions.

### Functional repertoire analyses

nAbs VH and VL sequence reads were manually curated and retrieved using CLC sequence viewer (Qiagen). Aberrant sequences were removed from the data set. Analyzed reads were saved in FASTA format and the repertoire analyses was performed using Cloanalyst (http://www.bu.edu/computationalimmunology/research/software/)^29,30^.

### Radar plot distribution of IGHV;IGHJ germlines

A radar plot was generated to display the distribution of IGHV;IGHJ germlines among the three nAbs groups: seronegative 2^nd^ dose, 3^rd^ dose and seropositive 2^nd^ dose. Each star in the radar represents a particular IGHV;IGHJ germline combination, alphabetically sorted. Germline abundance is represented as percentage of the total of each group. In addition, the log2 fold change for each combination of the three groups was calculated and represented as a concentric heatmap; the higher the fold change it is, the bigger is the first group with respect to the other, and vice versa. The figure was generated with R v4.1.1. and assembled with ggplot2 v3.3.5.

### Network plot of clonally expanded antibody families

To investigate the genetic similarity within and between lineages, a network map was built by representing each clonal family with a centroid and connecting centroids sharing a similar sequence. The centroid sequence was computed with Cloanalyst to represent the average CDRH3 sequence for each clonal family, and Hamming distance was calculated for each antibody CDRH3 sequence to represent the relationship within the clonal family. Levenshtein distance was calculated between each centroid representative of each clonal family to investigate the relationship between clonal families. Levenshtein distance was calculated with the R package stringdistm v0.9.8 (https://cran.r-project.org/web/packages/stringdist/index.html) and normalized between 0 and 1. A network graph was generated with the R package ggraph v2.0.5 (https://ggraph.data-imaginist.com/index.html) with Fruchterman-Reingold layout algorithm and the figure was assembled with ggplot2 v3.3.5. The size of the centroid is proportional to the number of antibodies belonging to the same clonal family, while the color of each node represents the antibody origin: light blue and dark blue for seronegative 2^nd^ dose and seronegative 3^rd^ dose, respectively.

### Statistical analysis

Statistical analysis was assessed with GraphPad Prism Version 8.0.2 (GraphPad Software, Inc., San Diego, CA). Nonparametric Mann-Whitney t test was used to evaluate statistical significance between the two groups analyzed in this study. Statistical significance was shown as * for values ≤ 0.05, ** for values ≤ 0.01, *** for values ≤ 0.001, and **** for values ≤ 0.0001.

## Acknowledgments

This work was funded by the European Research Council (ERC) advanced grant agreement number 787552 (vAMRes). This work was supported by a fundraising activity promoted by Unicoop Firenze, Coop Alleanza 3.0, Unicoop Tirreno, Coop Centro Italia, Coop Reno e Coop Amiatina. This publication was supported by the COVID-2020-12371817 project, which received funding from the Italian Ministry of Health. We would also like to acknowledge Dr. Jason McLellan, for kindly providing the S protein trimer, RBD, NTD and S2 constructs, Dr. Olivier Schwartz, for providing the B.1.617.2 (delta) SARS-CoV-2 variant, and Dr. Piet Maes for providing the B.1.1.529.1 (BA.1) and B.1.1.529.2 (BA.2) SARS-CoV-2 variants. We would like to thank the nurse staff of the operative unit of the department of Medical Sciences, Infectious and Tropical Diseases Unit, Siena University Hospital, Siena, Italy, and all the COVID-19 vaccinated donors for participating to this study.

## Author contributions

Conceived the study: E.A. and R.R.; Enrolled COVID-19 vaccinees: F.M., M.F., I.R. and M.T.; Performed PBMC isolation and single cell sorting: E.A and I.P.; Performed ELISAs and competition assays: I.P., V.A., G.A.; Recovered nAbs VH and VL and expressed antibodies: I.P. and N.M.; Recovered VH and VL sequences and performed the repertoire analyses: P.P., E.A. and G.M.; Produced and purified SARS-CoV-2 S protein constructs: E.P.; Performed neutralization assays in BSL3 facilities: E.A., G. Pie., G.Pic., M.L., L.B. and G.G.; Performed SARS-CoV-1 pseudotype neutralization assays: S.M. and L.D.; Supported day-by-day laboratory activities and management: C.D.S.; Manuscript writing: E.A. and R.R.; Final revision of the manuscript: E.A., I.P., G.Pie., G.Pic., V.A., G.A., P.P., N.M., E.P., L.B., G.G., M.L., S.M., L.D., C.D.S., M.F., I.R., M.T., F.M., C.S., R.D.F., E.M. and R.R.; Coordinated the project: E.A., C.S., E.M., R.D.F. and R.R.

## Additional information

**Correspondence and requests for materials** should be addressed to R.R.

## Data availability

Source data are provided with this paper. All data supporting the findings in this study are available within the article or can be obtained from the corresponding author upon request.

## EXTENDED DATA FIGURES

**Extended Data Fig. 1.**
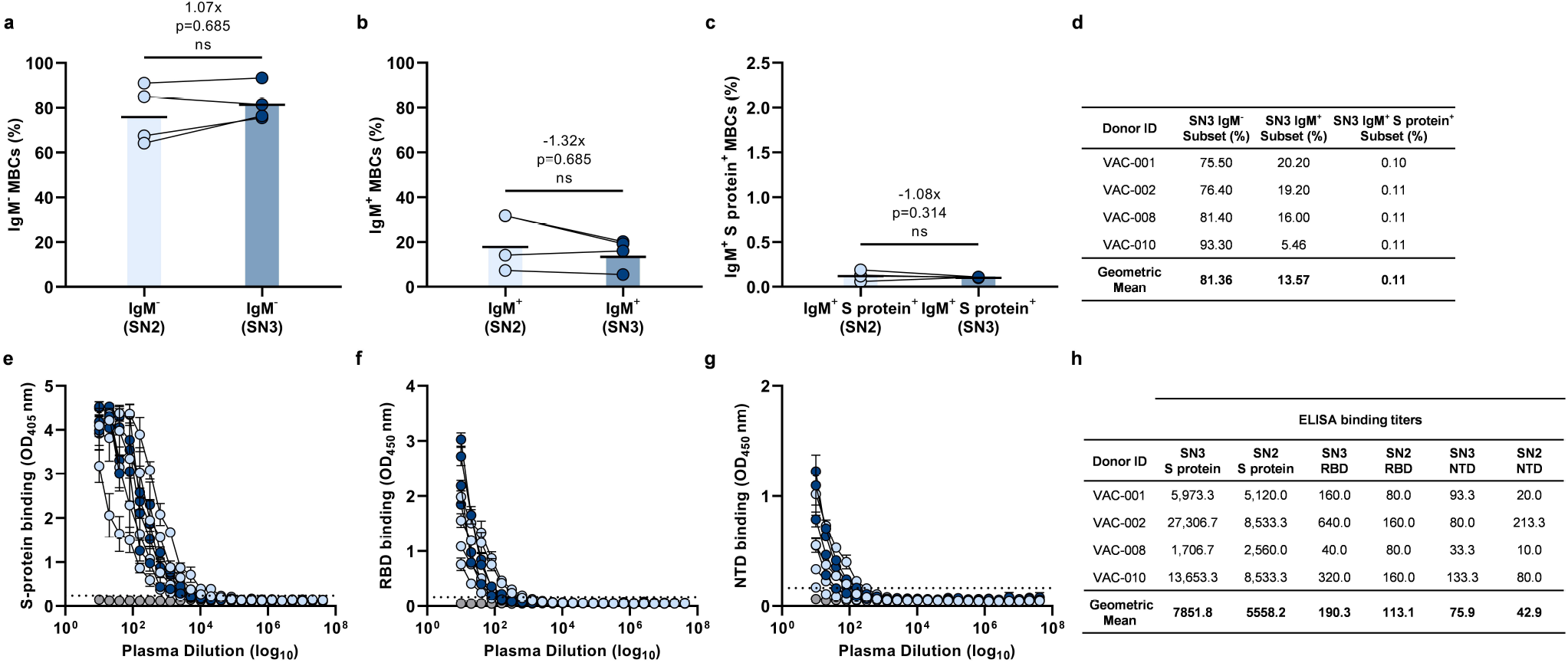
B cell frequencies and polyclonal response. **a-c**, The graph shows the frequency of CD19^+^CD27^+^IgD^-^IgM^-^ (**a**) and IgM^+^ (**b**), and CD19^+^CD27^+^IgD^-^IgM^+^ able to bind the SARS-CoV-2 S protein trimer (S protein^+^) (**c**) in SN2 and SN3 (*n* = 4/group). Black line and bars denote the geometric mean. **d**, The table summarizes the frequencies of the B cell populations for the SN3 group. **e-g**, Graphs show the ability of plasma samples from SN2 and SN3 to bind the S protein trimer (**e**), RBD (**f**) and NTD (**g**). Mean and standard deviation are denoted on each graph. Technical triplicates were performed for each experiment. **h**, The table summarizes the binding titers of each COVID-19 vaccinee and the geometric mean for SN2 and SN3. A nonparametric Mann–Whitney t test was used to evaluate statistical significances between groups. Two-tailed p-value significances are shown as *p < 0.05, **p < 0.01, ***p < 0.001, and ****p < 0.0001.

**Extended Data Fig. 2.**
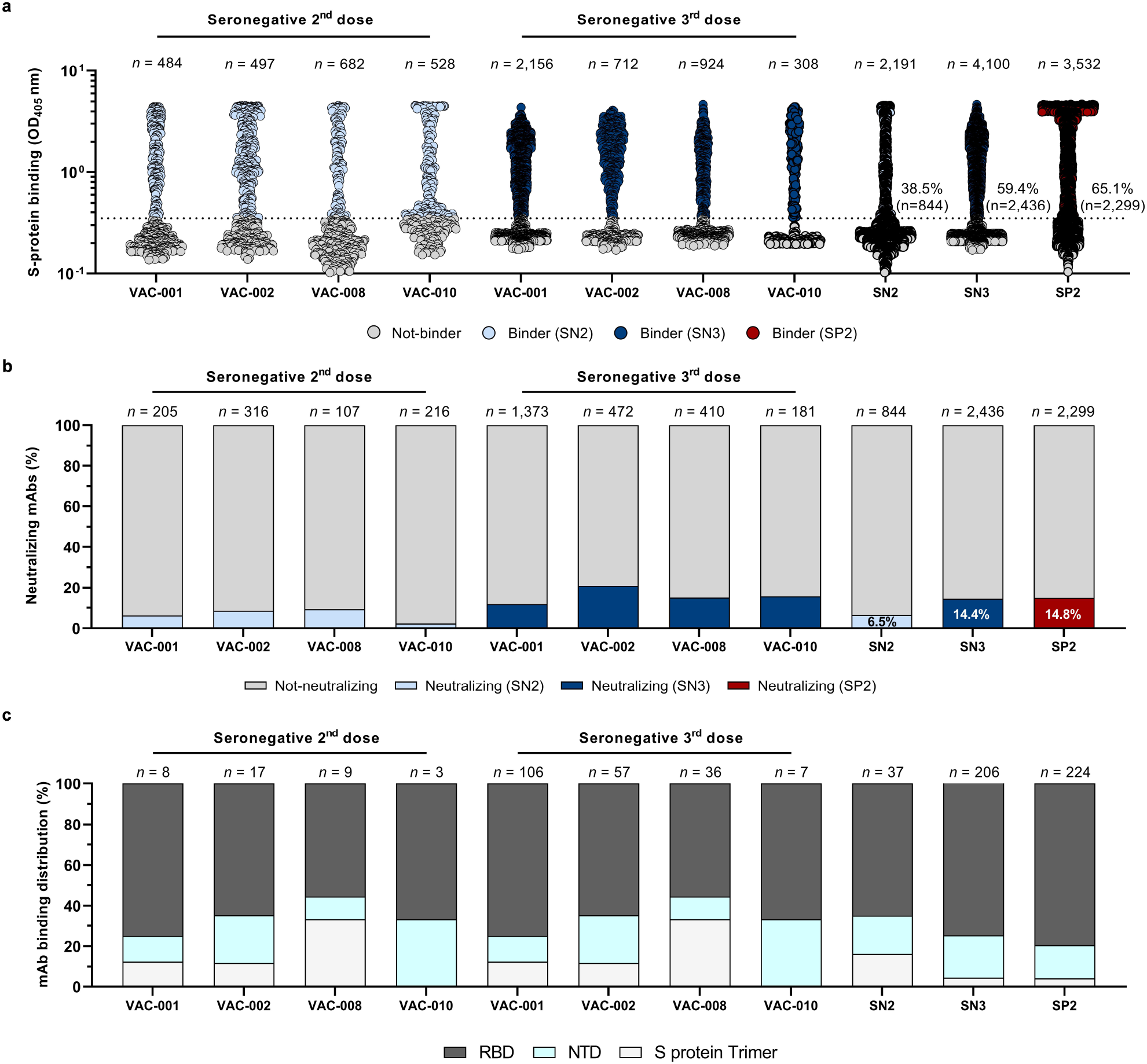
Identification of SARS-CoV-2 S protein-specific nAbs. **a**, The graph shows supernatants tested for binding to the Wuhan SARS-CoV-2 S protein antigen. Threshold of positivity has been set as two times the value of the blank (dotted line). Light blue, dark blue and red dots represent mAbs that bind to the S protein for SN2 and SN3, and SP2 respectively. Light gray dots represent mAbs that did not bind the S protein. **b**, The bar graph shows the percentage of not-neutralizing (gray), neutralizing mAbs from SN2 (light blue), neutralizing mAbs for SN3 (dark blue) and neutralizing mAbs for SP2 (red). The total number (*n*) of antibodies tested per individual is shown on top of each bar. **c**, The graph shows the percentage of nAbs that bind specifically the RBD (dark gray), the NTD (cyan) or that did not bind single domains but recognized exclusively the S protein in its trimetric conformation (light gray). The number (*n*) of tested nAbs per donor is reported on top of each bar. Technical duplicates were performed for each experiment.

**Extended Data Fig. 3.**
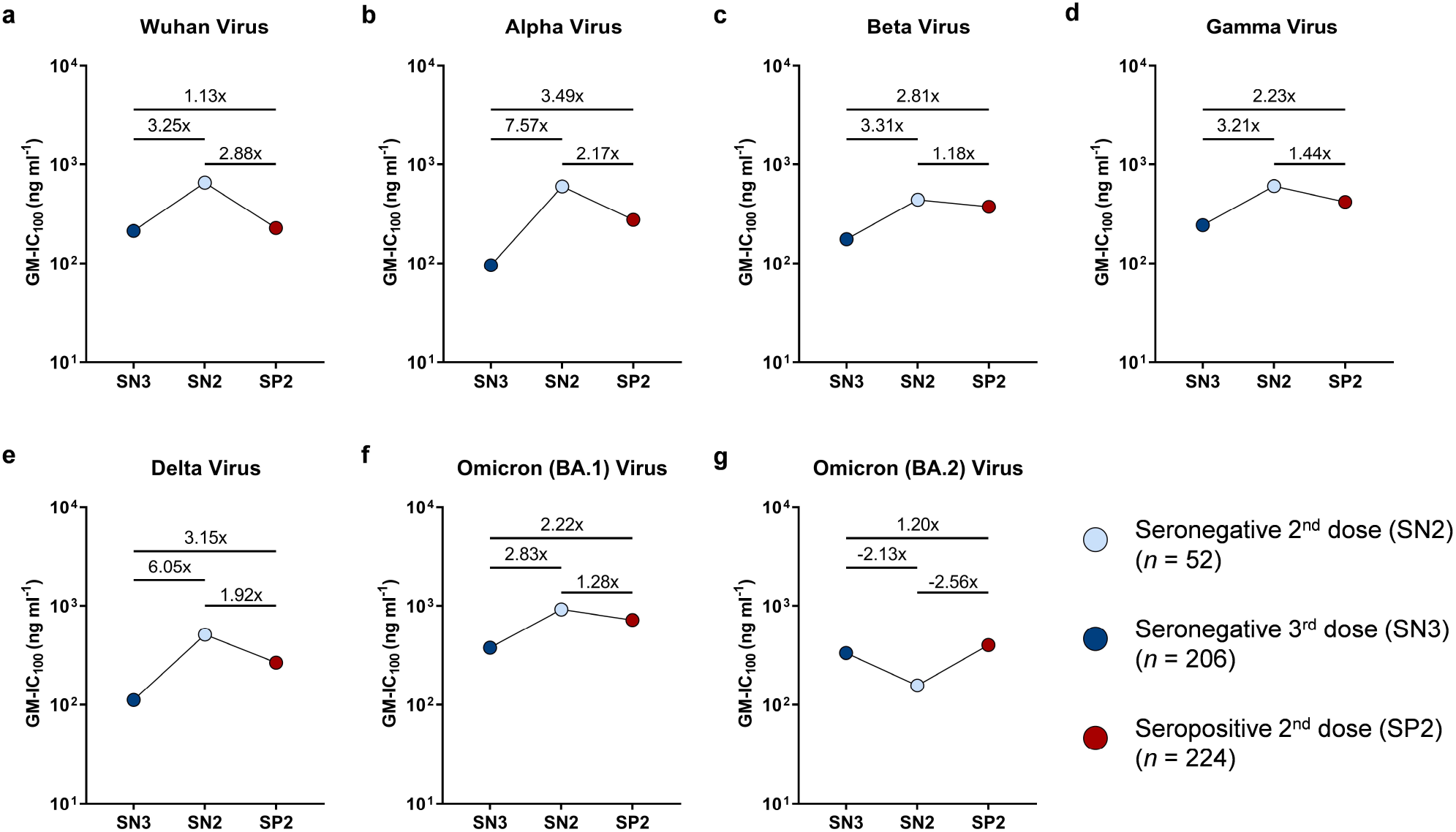
Fold-change neutralization potency against SARS-CoV-2 VoCs. **a-g**, The graphs show the fold-change neutralization potency shown as GM-IC_100_ (ng ml^-1^) among SN2 (light blue), SN3 (dark blue) and SP2 (red). Fold-changes are denoted on each graph.

**Extended Data Fig. 4.**
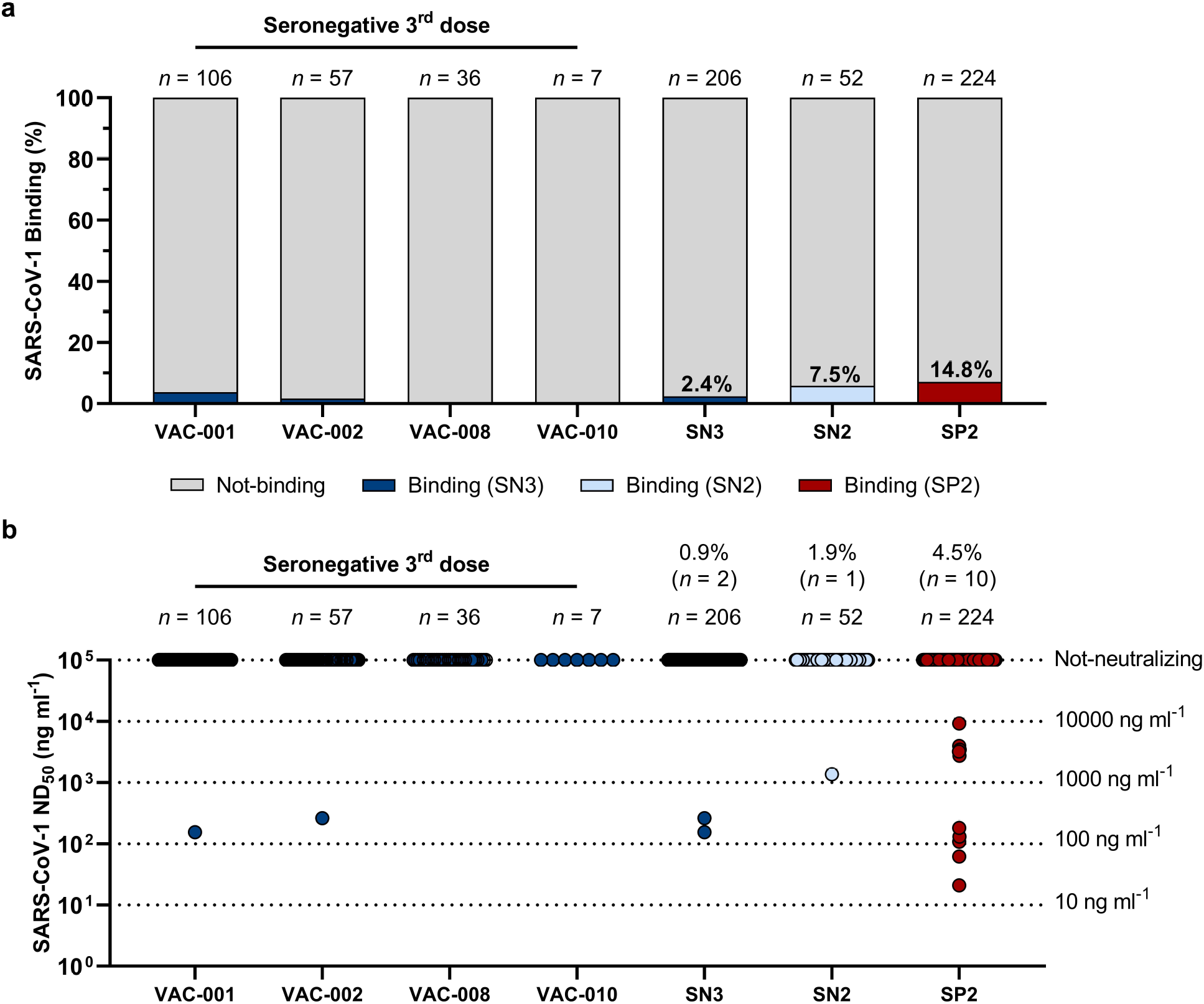
Functional characterization of SARS-CoV-1 nAbs. **a**, The bar graph shows the percentage of not-binding antibodies (grey), and SARS-CoV-1 binding nAbs for SN3 (dark blue), SN2 (light blue) and SP2 (red). The total number (*n*) of antibodies tested per individual is shown on the top of each bar. **b**, Dot chart shows the neutralization potency, reported as 50% neutralizing dilution (ND_50_ ng ml^−1^), of nAbs isolated from SN3 (dark blue), SN2 (light blue) and SP2 (red). The number and percentage of nAbs from individuals who were seronegative and seropositive and neutralization ND_50_ (ng ml^−1^) ranges (black dotted lines) are denoted on the graph.

**Extended Data Fig. 5.**
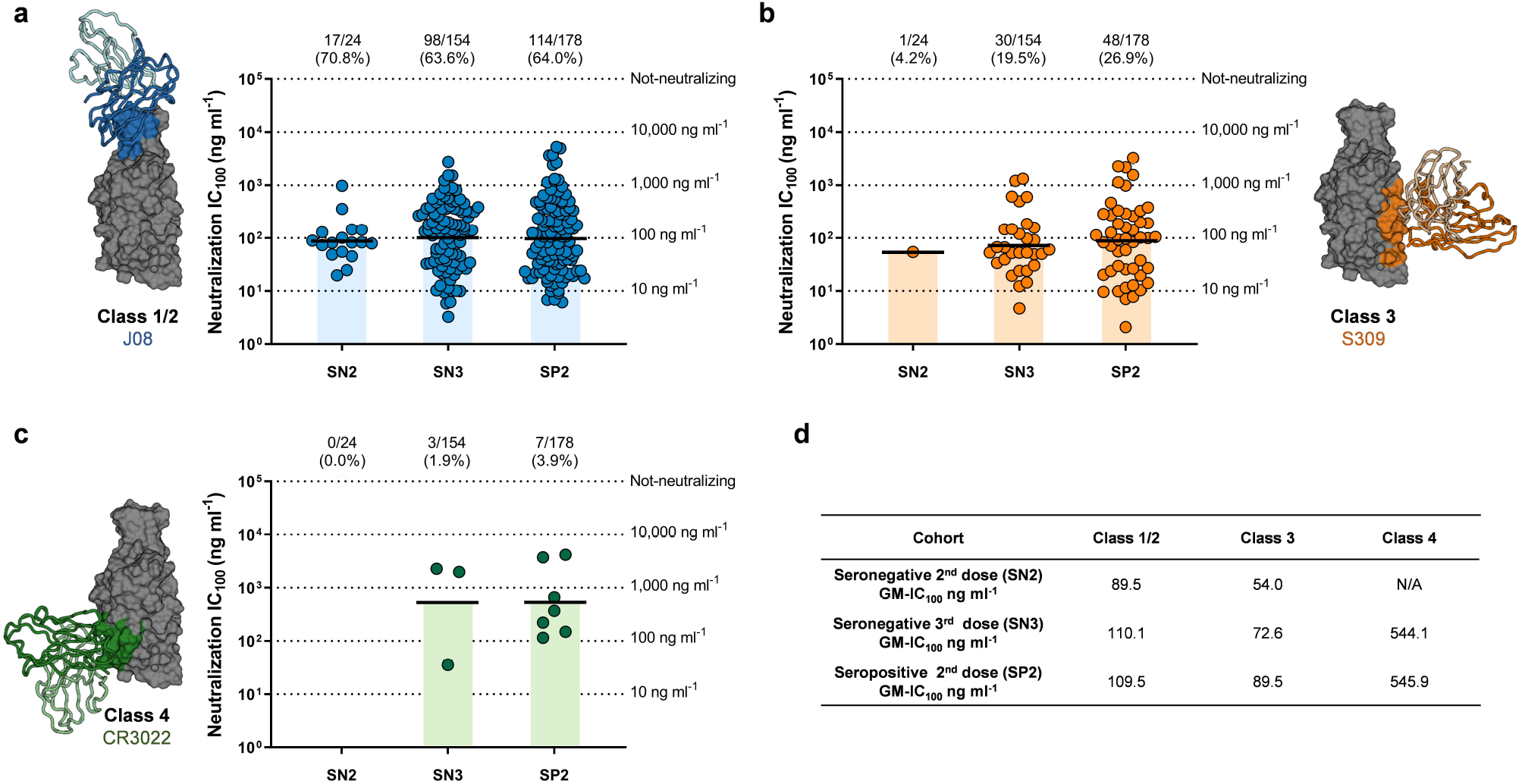
Epitope mapping of RBD-targeting nAbs. **a-c**, Dot charts show the distribution of Class 1/2 (**a**), Class 3 (**b**) and Class 4 (**c**) nAbs against the original SARS-CoV-2 virus first detected in Wuhan for nAbs isolated from SN2, SN3 and SP2. The number and percentage of nAbs and neutralization IC_100_ geometric mean (black lines, light blue, orange and green bars) are denoted on each graph. **d**, the table summarizes the IC_100_ geometric mean of nAbs against Wuhan for all tested groups.

**Extended Data Fig. 6.**
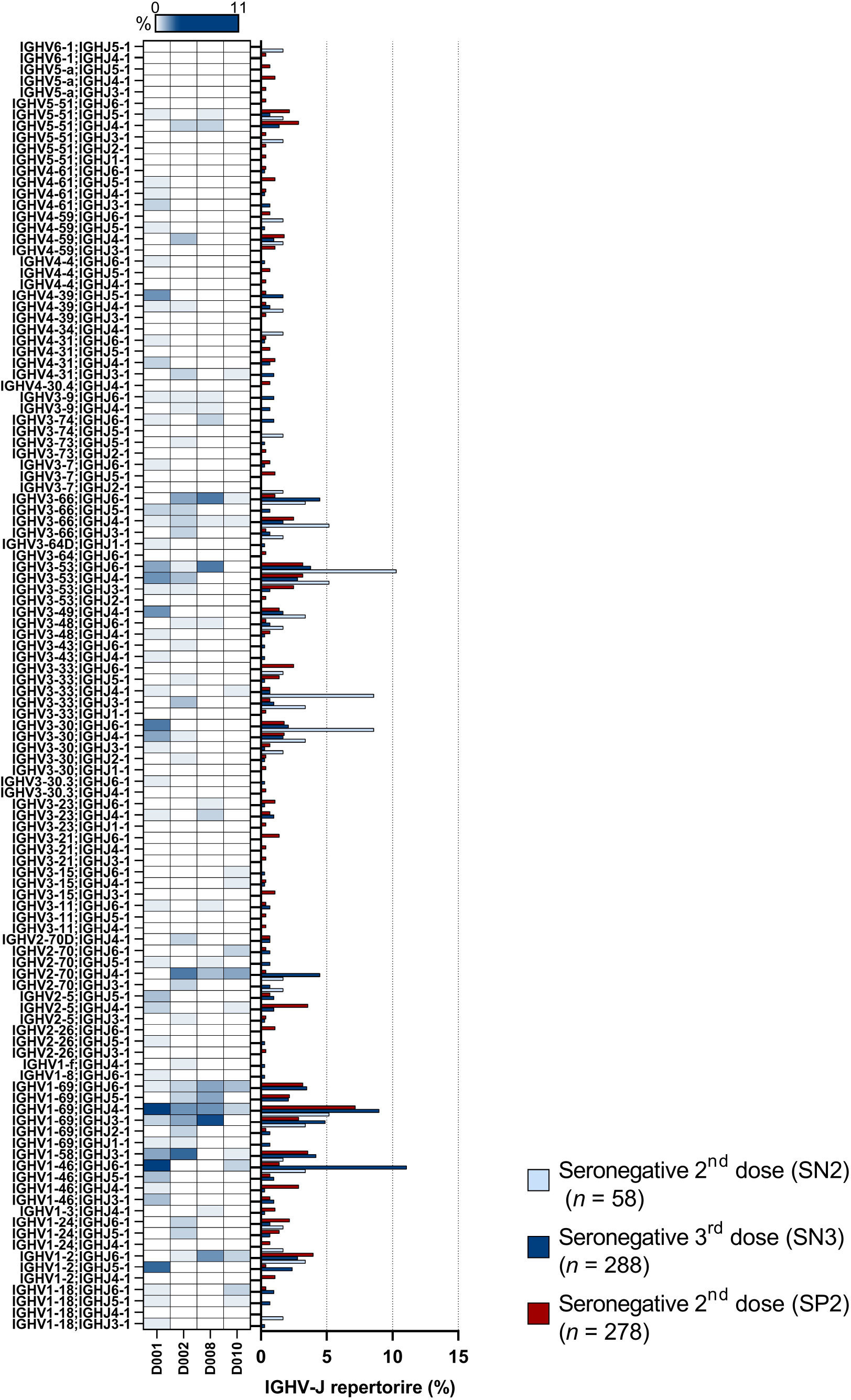
Antibody germline distribution. The graph shows the IGHV;IGHJ rearrangement frequencies among SN2 (light blue), SN3 (dark blue) and SP2 (red) dose vaccinees (right panel), and the frequency within SN3 subjects (left panel).

## EXTENDED DATA TABLES

**Extended Data Table 1.** Clinical details of COVID-19 vaccinees.

| Subject ID | Gender | Age | First Dose (dd/mm/yy) | Second Dose (dd/mm/yy) | Third Dose (dd/mm/yy) | Vaccine | Days from Second to Third Dose | Blood Collection (dd/mm/yy) | Days from Third Dose to Blood Collection |
| --- | --- | --- | --- | --- | --- | --- | --- | --- | --- |
| VAC-001 | M | 39 | 27/12/2020 | 18/01/2021 | 28/10/2021 | BNT162b2 | 283 | 13/01/2022 | 77 |
| VAC-002 | F | 39 | 01/01/2021 | 22/01/2021 | 08/11/2021 | BNT162b2 | 290 | 13/01/2022 | 66 |
| VAC-008 | M | 44 | 03/01/2021 | 24/01/2021 | 23/11/2021 | BNT162b2 | 303 | 12/01/2022 | 50 |
| VAC-010 | F | 52 | 18/02/2021 | 11/03/2021 | 09/12/2021 | mRNA-1273 | 277 | 20/01/2022 | 42 |

**Extended Data Table 2.** Summary of sorted B cells and neutralizing antibodies against SARS-CoV-2 and VoCs.

| Subject | S protein<sup>+</sup><br>MBCs Sorted | S protein<sup>+</sup><br>mAbs (%) | Wuhan -<br>Neutralizing<br>antibodies (%) | Expressed -<br>Neutralizing<br>antibodies<br>(%) | Expressed -<br>Neutralizing<br>antibodies (%)<br>Alpha | Expressed -<br>Neutralizing<br>antibodies (%)<br>Beta | Expressed -<br>Neutralizing<br>antibodies (%)<br>Gamma | Expressed -<br>Neutralizing<br>antibodies (%)<br>Delta | Expressed -<br>Neutralizing<br>antibodies (%)<br>Omicron BA.1 | Expressed -<br>Neutralizing<br>antibodies (%)<br>Omicron BA.2 |
| --- | --- | --- | --- | --- | --- | --- | --- | --- | --- | --- |
| VAC-001 | 2,156 | 1,373 (63.7) | 162 (11.8) | 106 (65.4) | 68 (64.2) | 36 (34.0) | 30 (28.3) | 55 (51.9) | 22 (20.8) | 24 (22.6) |
| VAC-002 | 712 | 472 (66.3) | 99 (21.0) | 57 (57.6) | 54 (94.7) | 34 (59.6) | 35 (61.4) | 42 (73.7) | 15 (26.3) | 17 (29.8) |
| VAC-008 | 924 | 410 (44.4) | 61 (14.9) | 36 (59.0) | 31 (86.1) | 14 (38.9) | 14 (38.9) | 25 (69.4) | 7 (19.4) | 9 (25.0) |
| VAC-010 | 308 | 181 (58.8) | 28 (15.5) | 7 (25.0) | 6 (85.7) | 2 (28.6) | 2 (28.6) | 4 (57.1) | 1 (14.3) | 2 (28.6) |
| Total | 4,100 | 2,436 (59.4) | 350 (14.4) | 206 (51.7) | 159 (77.2) | 86 (41.7) | 81 (39.3) | 126 (61.2) | 45 (21.8) | 52 (25.2) |

